# MetalPrognosis: a Biological Language Model-based Approach for Disease-Associated Mutations in Metal-Binding Site prediction

**DOI:** 10.1101/2023.11.01.565246

**Authors:** Runchang Jia, Zhijie He, Cong Wang, Xudong Guo, Fuyi Li

**Author notes:** Fuyi Li is the corresponding author.

## Abstract

Protein-metal ion interactions play a central role in the onset of numerous diseases. When amino acid changes lead to missense mutations in metal-binding sites, the disrupted interaction with metal ions can compromise protein function, potentially causing severe human ailments. Identifying these disease-associated mutation sites within metal-binding regions is paramount for understanding protein function and fostering innovative drug development. While some computational methods aim to tackle this challenge, they often fall short in accuracy, commonly due to manual feature extraction and the absence of structural data. We introduce MetalPrognosis, an innovative, alignment-free solution that predicts disease-associated mutations within metal-binding sites of metalloproteins with heightened precision. Rather than relying on manual feature extraction, MetalPrognosis employs sliding window sequences as input, extracting deep semantic insights from pre-trained protein language models. These insights are then incorporated into a convolutional neural network, facilitating the derivation of intricate features. Comparative evaluations show MetalPrognosis outperforms leading methodologies like MCCNN and PolyPhen-2 across various metalloprotein test sets. Furthermore, an ablation study reiterates the effectiveness of our model architecture. To facilitate public use, we have made the datasets, source codes, and trained models for MetalPrognosis online available at http://metalprognosis.unimelb-biotools.cloud.edu.au/.

## 1 Introduction

Research indicates that metal ions interact with over half of all proteins, reinforcing their structures and enabling catalytic functions [1]. Such interactions induce protein conformation alterations, affecting enzyme catalysis, homeostasis regulation, and numerous physiological functions. These interactions extend beyond mere structural stabilization; they profoundly impact the proteins’ three-dimensional form, stability, and overall biological functionality [2,3]. At metal binding sites, amino acids coordinate directly with metals, determining the wide-ranging functionalities of metalloproteins. These roles encompass structural maintenance, dioxygen binding, electron transfer, molecular recognition, gene transcription, and regulation of expression [4,5,6]. For example, metal ion (*Fe*^3+^) binding to hemoglobin is vital for oxygen transport in blood [7,8]. Metal ions also underpin myriad biological roles, such as the regulation of muscle strength by intracellular calcium concentration [9] and the interactions amino acids establish with metals via mechanisms like hydrogen bonding, hydrophobic interactions, and van der Waals forces [10].

However, single nucleotide point mutations can cause amino acid coding changes, leading to missense mutations [11]. When these mutations manifest at metal-binding sites, they can obstruct the protein-metal ion interaction, potentially undermining protein function. Such disruptions are linked to severe human conditions like Epidermolysis bullosa and sickle-cell disease [12,13]. Initial methods like SIFT, which primarily depend on evolutionary data, were developed to detect these mutations. SIFT is heavily reliant on PSI-BLAST’s sequence alignment data, suggesting that mutations in highly conserved protein sequences frequently result in harmful outcomes [14,15,16].

In the recent past, machine learning and data mining techniques have been leveraged to probe pathogenic missense mutations in the human genome [17,18]. Techniques like CHASM use random forest and SVM classifiers to distinguish between driver mutations and synthetically generated passenger mutations in cancer-specific somatic cells [19]. Another method, PolyPhen-2, amalgamates both sequence and structural data. It collates information from sequence annotations, multiple sequence alignments, and existing 3D structures, utilizing a naive Bayesian classification approach [20]. Despite their effectiveness, these techniques depend heavily on conventional machine learning algorithms, demanding significant computational resources and time, particularly when gleaning evolutionary data via multiple sequence alignment (MSA) [21]. In scenarios where protein sequences lack extensive homologous families, this becomes a limitation. On the other hand, deep learning can auto-extract features, improving model generalization [22,23,24,25].

The advent of advanced computational capabilities has seen deep learning make significant strides in predicting pathogenic missense mutations [26,27,28,29,30]. For instance, MCCNN incorporates omics data to pinpoint missense mutations, using energy-based affinity grids and physicochemical attributes as input for a 3D convolutional neural network [31]. However, there is ample scope for enhancement. For instance, the grid length in MCCNN is merely 15Å, limiting its extraction capabilities. MVP, another model, utilizes six interrelated features for predictions and operates on residual neural networks [32]. These scoring matrices for amino acid substitution, such as BLOSUM62 and PAM250, generate fixed evolutionary information patterns, potentially failing to capture dynamic high-dimensional features based on varying metal physiological environments.

The emergence of transformers and unsupervised learning on vast protein datasets has further propelled the field, especially concerning tasks like protein tertiary structure prediction and protein-protein interaction [33,34,35]. LMetal-Site, for instance, utilizes a pre-trained language model to swiftly generate data and predict protein metal ion binding sites, outclassing all static sequence feature-centric methods [36]. Dynamic embedding’s paramount advantage is its ability to extract diverse semantic information based on various metal protein site contexts.

In this research, we introduce MetalPrognosis, a pioneering alignment-free deep learning approach tailored for such predictions. Designed to capture the long-term dependency information between amino acids, it extracts dynamic semantic features from the ESM pre-trained protein language model. These are subsequently processed by a convolutional neural network to extract abstract features. Notably, MetalPrognosis consistently surpasses competitors, such as MCNN, on multiple evaluation metrics, including specificity and accuracy. Additionally, MetalPrognosis plays a pivotal role in identifying pathogenic missense mutation sites in human metalloproteins, highlighting its significant potential in disease diagnosis. To facilitate public use, We provide an online prediction platform at http://metalprognosis.unimelb-biotools.cloud.edu.au/.

## 2 Material and Methods

### 2.1 Data collection and preprocessing

The initial phase involved data collection and cleaning (Figure 1A). To assess MetalPrognosis’ efficacy, we utilized the dataset from MCCNN, available in the supplementary information section. This dataset amalgamated omics data from four databases to pinpoint missense mutations at the metal-binding sites of proteins. We began by downloading 169,123 PDB structures of recognized metal-binding sites from MetalPDB https://metalpdb.cerm.unifi.it/. Of these, 22,307 protein structures were shortlisted based on two criteria: the spatial distance between the protein structure’s alpha carbon atom and the specified metal ion being less than 10Å, and their origin being human species. From three human disease-associated databases—ClinVar, Uniprot Hamsavar, and CancerResource2—we extracted mutations amounting to 478, 442, and 509, respectively. Furthermore, 261 distinctive benign missense mutations situated in the metal-binding sites of proteins were gathered. For ID mapping, we navigated to the UniProt website https://www.uniprot.org/, inputted the previously collected PDB IDs, and awaited the results. Upon completion, we had the option to download the amino acid sequence data in either FASTA or TSV format.

**Fig. 1.**
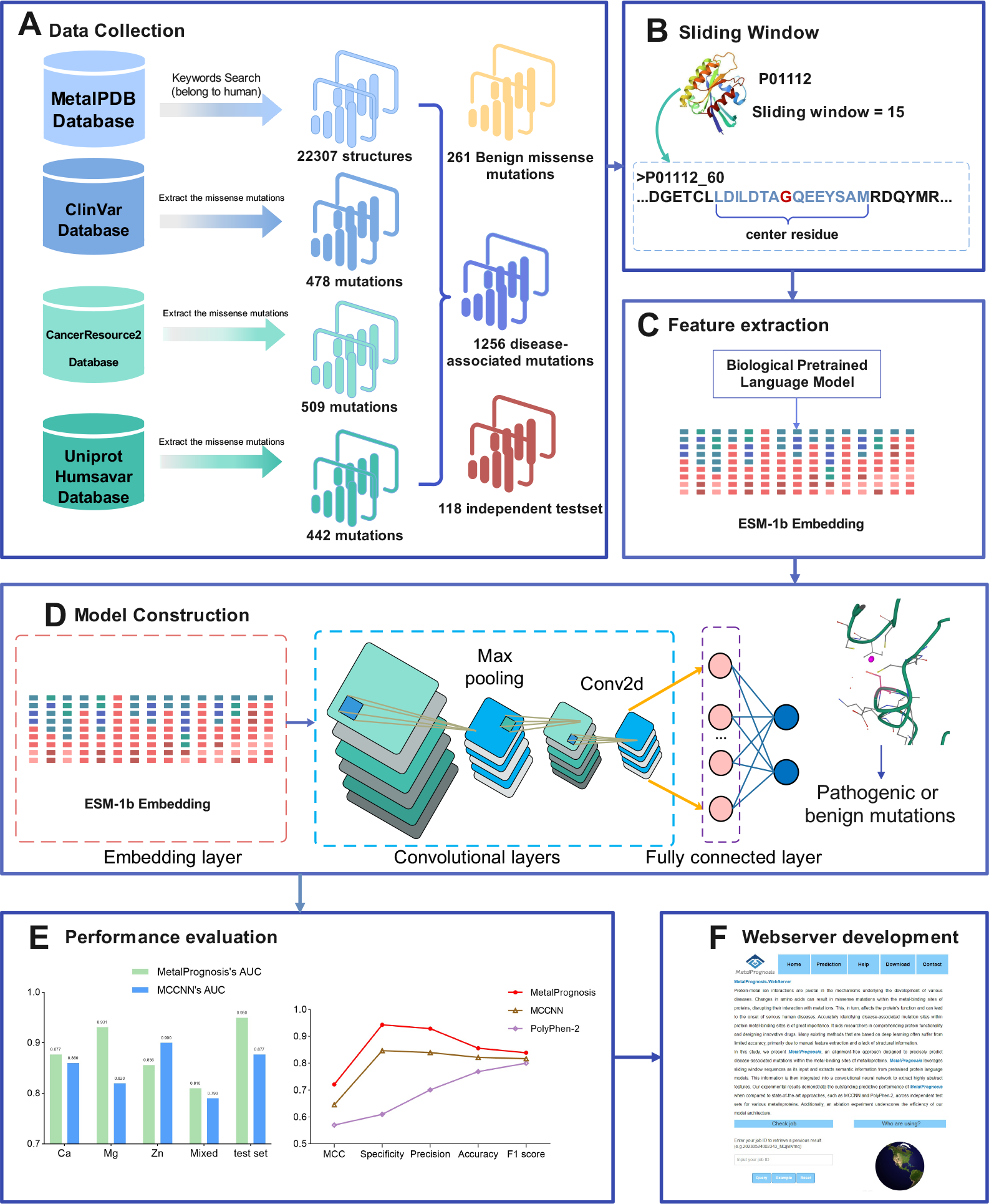
The overall framework of MetalPrognosis. This flowchart consists of 6 parts, including **A**. Data collection. **B**. Use a sliding window to extract subsequences centred at the mutation site. **C**. Extracting semantic features from protein pre-trained language model. **D**. Model construction, the architecture of the proposed model. **E**. Performance evaluation, results of comparison experiments. **F**. Webserver development.

We utilized disease-associated mutations as positive samples and benign mutations as negative samples for training the deep learning model. The independent test sets comprised 59 disease-associated mutations and an equal number of benign mutations, each sourced from distinct metal binding sites. By including diverse disease-associated mutations from various metal binding sites, such as *Ca*^2+^, *Mg*^2+^, and *Zn*^2+^ ions, our objective was to ensure a thorough and unbiased comparison of MetalPrognosis with pre-existing methodologies on these independent test sets. Subsequently, the benchmark dataset was partitioned into training and validation subsets. To provide specifics: the counts for zinc, calcium, magnesium binding sites, and benign mutations stood at 447, 334, 171, and 261, respectively. The details of the benchmark dataset are provided in Table 1, facilitating further analysis and juxtaposition.

**Table 1:**
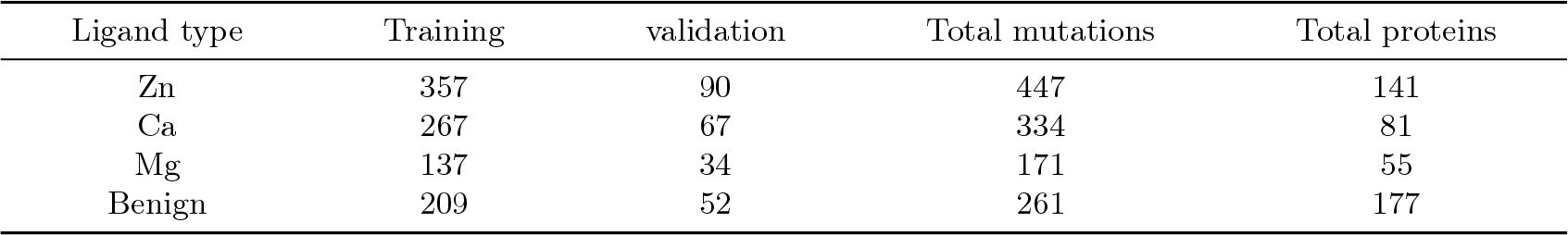
Statistics on disease-associated mutations of three metal binding sites and benign mutations.

| Ligand type | Training | validation | Total mutations | Total proteins |
| --- | --- | --- | --- | --- |
| Zn | 357 | 90 | 447 | 141 |
| Ca | 267 | 67 | 334 | 81 |
| Mg | 137 | 34 | 171 | 55 |
| Benign | 209 | 52 | 261 | 177 |

### 2.2 The overall framework of MetalPrognosis

Figure 1 illustrates the overall framework of MetalPrognosis. As can be seen, it is built sequentially from 6 submodules, including (A)Data collection, (B) Sliding window module, (C) Feature extracting module, (D) Deep learning model construction, (E) Performance evaluation, and (F) Webserver development.

To begin with, we collected the dataset from four different databases, finding missense mutations that occur in the metal-binding sites of proteins. There are 1256 disease-associated mutations and 261 benign mutations for training the model. 118 mutations at the metal binding site were used as independent test sets to verify the generalization ability of the model. In the second stage, we use a sliding window to extract subsequences centered at the mutation site. In the third stage, amino acid subsequences were input into the protein pretrained language model to extract semantic features. The next step is model construction and training. We construct a two-layer convolutional neural network for high-dimensional feature extraction. The next step is performance evaluation. We evaluated and compared the predictive performance of MetalPrognosis with state-of-the-art methods on an independent test set. The last step is webserver development, and we developed an online web server that is convenient for the public to use.

### Sliding window

Full-length protein sequences were not used as model input due to the excessive lengths of some metal-binding proteins, which could dilute the focus on mutation sites. The binding of a protein to a metal ion involves not just the binding residue but also the adjacent residues, which influence the interaction. To capture context and distant interactions of the mutation site, we employed a sliding window method to segment amino acid sequence fragments (Figure 1B). The chosen window size is crucial as it determines the depth of context-dependent information from the amino acid sequence [37,38]. While a larger window can yield richer context, it might compromise the model’s predictive efficiency [39]. To maintain the feature matrix’s width, edge samples for mutation sites at sequence extremes were padded with Alanine. Prior research indicated optimal results with seven amino acids both upstream and downstream of the mutation site, thus setting our sliding window size to 15 [40]. Fragments centered on disease-related metal-binding sites were labeled as positive samples, whereas those on benign mutation sites were labeled negative.

### Pre-trained language model representation

Conventional embedding methods like AAC (Amino Acid Composition) employ a fixed coding scheme for each amino acid. This approach fails to capture the dynamic semantic nuances of amino acid sequences based on their context [41]. To overcome this limitation, we leveraged the ESM (Evolutionary Scale Model) — a protein pretrained language model — to extract dynamic global context information from amino acid sequences (Figure 1C). Specifically, our choice was the esm1b t33 650M UR50S model [42], an unsupervised protein language model inspired by the Bidirectional Encoder Representations from Transformers (BERT) design. Through self-supervised training on a vast corpus of unlabeled protein data, it employs techniques like masking language modeling and next-sentence prediction. This allows the model to capture intricate details about semantic nuances and structural attributes in protein sequences, placing it at the forefront of multiple protein-related tasks.

The strength of unsupervised models like ESM-1b lies in their ability to harness the wealth of sequence data in protein databases without the need for manual annotations or explicit homologies. ESM-1b’s training foundation is the UniRef protein sequence database [43], involving uniform sampling from 43 million UniRef50 training clusters derived from 138 million UniRef90 sequences. This model employs a BERT-esque learning approach, focusing on semantic and structural features within protein sequences. With its 33 layers, 650M parameters, and a 1280-dimensional embedding for each amino acid, the model encompasses information related to secondary structure, physicochemical properties, homology, and more. Evidence suggests that ESM-1b can deduce myriad facets of a protein’s 3D structure and function purely from its sequence, outperforming in several protein-centric tasks, such as determining secondary structures, pinpointing protein binding sites, and gauging mutational impacts [44,45]. Additionally, we evaluated another protein language model, ProtTrans [46]. Known as ProtBert-BFD, it undergoes unsupervised training on the BFD100 dataset, features 30 layers, a hidden layer size of 1024, a mask probability of 15%, and a total of 420M parameters.

### The architecture of the deep learning model

MetalPrognosis treats this task as a binary classification task: determining, for a given metalloprotein binding site in a human species, whether it corresponds to a disease-related mutation site or a benign mutation site. As illustrated in Figure 1D, MetalPrognosis operates as an integrated deep learning model utilizing convolutional neural networks, eliminating the need for intricate manual feature design. The embedded feature matrix we extracted stands at a dimensionality of 1280, serving as the CNN model’s feature map. Following this, the convolutional layer extracts abstract feature maps, succeeded by a max-pooling layer to downsize the feature maps’ dimensionality, effectively halving the original resolution and reducing the model’s parameter count. To enhance the model’s generalizability, a batch normalization layer is incorporated post-convolution. In the fully connected segment, a dropout rate of 0.3 is implemented to mitigate overfitting. Ultimately, the fully connected network merges with the Sigmoid classifier to produce binary classification outcomes. Additionally, MetalPrognosis was constructed using PyTorch and honed on a high-capacity Linux server. Comprehensive implementation details, including hyperparameter configurations, are elaborated upon in the supplementary material.

## 3 Results and discussion

### 3.1 Performance evaluation on the training dataset

In this study, due to a sufficient number of samples for zinc, calcium, and magnesium ion disease-associated binding sites, we focused on these three metal types. A 5-fold cross-validation strategy was employed to train MetalPrognosis, and its performance was assessed using accuracy, precision, recall, F1-score, AUPRC, MCC, and AUC. Detailed descriptions of these evaluation metrics are provided in the supplementary material.

For both zinc and calcium ions, we used all available mutation sites as positive samples and combined them with benign mutation sites for further feature extraction and model training. However, the dataset for magnesium ions was smaller, prompting us to downsample by randomly selecting 200 benign mutation sites to achieve balanced data, akin to the MCCNN approach. To assess the model’s generalization ability, we created a new dataset comprising various metal ions. This was done by randomly selecting 100 samples from disease-related mutation sites of each of the three metal ions, yielding a mixed positive dataset of 300 samples. This mixed set was then combined with benign mutation data for model training, ensuring a balanced ratio between positive and negative samples. This balance helps mitigate data bias and enhance model accuracy[47]. The amino acid distributions at different metal binding sites are detailed in the supplementary material.

Table 2 presents the performance results for four datasets. The zinc ion protein dataset, being the largest, achieved an accuracy of 0.931 and an AUC of 0.856. The calcium ion protein dataset had a slightly lower accuracy compared to zinc but boasted a marginally higher AUC of 0.877. The magnesium ion dataset, being the smallest, achieved an F1 of 0.818, AUPRC of 0.916, MCC of 0.691, and AUC of 0.931. However, the mixed dataset’s MCC and AUC were 0.485 and 0.81, respectively, lower than the results from individual metal datasets. This suggests varying physiological environments in metal binding pockets might lead to a decreased AUC.

**Table 2:** The main indicators of MetalPrognosis on different metal ion data set.

| Metal | Accuracy | Precision | Recall | F1-score | AUPRC | MCC | AUC |
| --- | --- | --- | --- | --- | --- | --- | --- |
| Ca | 0.851 | 0.666 | 0.571 | 0.615 | 0.713 | 0.526 | 0.877 |
| Mg | 0.840 | 0.750 | 0.900 | 0.818 | 0.916 | 0.691 | 0.931 |
| Zn | 0.931 | 0.610 | 0.701 | 0.769 | 0.801 | 0.570 | 0.856 |
| Mixed | 0.845 | 0.731 | 0.845 | 0.784 | 0.803 | 0.485 | 0.810 |

Figure 2A demonstrates that our model’s AUC outperformed MCCNN across three 5-fold cross-validation datasets and an independent test set, with the sole exception being the zinc ion dataset. Relative to MCCNN, our model’s AUC superiority was by margins of 0.111, 0.017, 0.02, and 0.073 for magnesium, calcium, the mixed dataset, and the independent test set, respectively. However, for the zinc ion dataset, while our model had an AUC of 0.856, MCCNN reached 0.90. This discrepancy might stem from MCCNN’s superior extraction of energy-based affinity grid spatial features from zinc ions or differing cross-validation strategies affecting dataset splits.

**Fig. 2.**
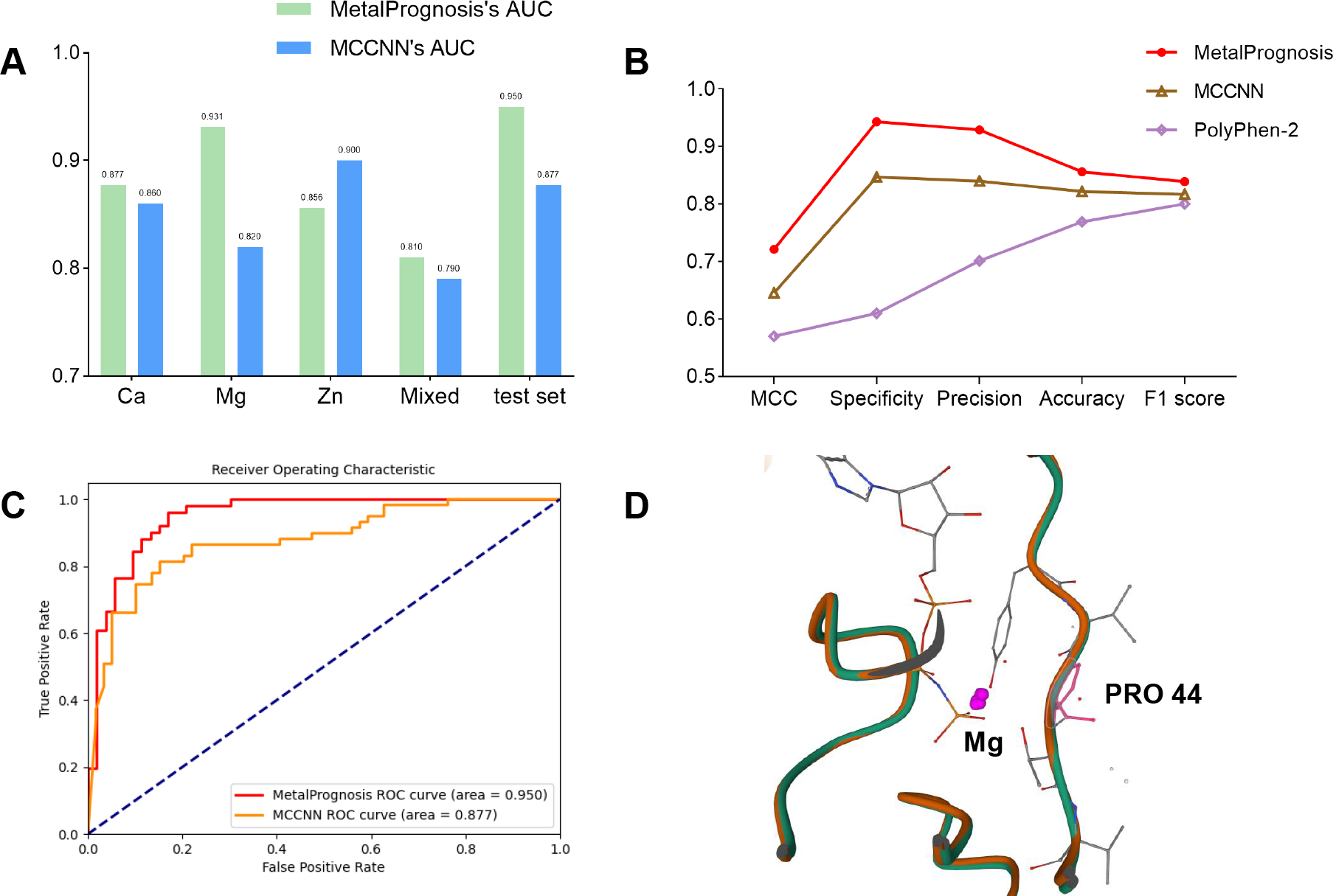
The performance comparison chart. **A**. Comparison of AUC performance on different metal ion datasets. **B**. The performance comparison between MetalPrognosis and other state-of-the-art approaches. **C**. The AUC performance comparison between MetalPrognosis and MCCNN. **D**. Benign mutation predicted by MetalPrognosis(Uniprot ID: Q92730).

### 3.2 Ablation test

To ascertain the efficacy of harnessing dynamic sequence context embedding from pre-trained protein language models, we undertook ablation experiments. These aimed to assess the sequence features extracted by various pre-trained language models and the influence of attention mechanisms on model performance. Initially, we juxtaposed the MCC and AUC performance metrics on three metal ion datasets between the ESM and ProtTrans models while keeping the neural network architecture consistent. As depicted in Table 3, the ESM model consistently outshines ProtTrans in both metrics. This discrepancy can be attributed to their unique training datasets— with ESM and ProtTrans being unsupervised trained on UniRef and BFD100 protein sequence datasets, respectively— and their distinct architectures and parameter configurations. We hypothesize that the masking protein modeling utilized in ESM pretraining enables its encoder to discern intricate relationships and patterns within protein sequences more effectively.

**Table 3:** Results of the ablation analysis of the MetalPrognosis.

| Feature | Ca MCC | Ca AUC | Zn MCC | Zn AUC | Mg MCC | Mg AUC | Mixed MCC | Mixed AUC |
| --- | --- | --- | --- | --- | --- | --- | --- | --- |
| ProtTrans | 0.519 | 0.796 | 0.493 | 0.826 | 0.659 | 0.900 | 0.467 | 0.781 |
| ESM+ ProtTrans | 0.515 | 0.833 | 0.548 | 0.855 | 0.685 | **0.937** | 0.479 | 0.726 |
| ESM+Attention | 0.480 | 0.815 | 0.524 | 0.828 | 0.676 | 0.915 | **0.487** | 0.803 |
| ESM(MetalPrognosis) | **0.526** | **0.877** | **0.570** | **0.856** | **0.691** | 0.931 | 0.485 | **0.810** |

To further our understanding, we combined two distinct dynamic sequence context embeddings, using them as the feature maps for the convolutional neural network to gauge any potential performance impact. Based on the results from a 5-fold cross-validation test, the AUC for the magnesium ion dataset registered at 0.937, a minor improvement over MetalPrognosis’s 0.931. However, for the remaining datasets, ESM’s performance surpassed that of the model post-feature concatenation. This suggests that embeddings derived from different extensive language models might possess overlapping features, and their efficacy varies when applied to specific downstream tasks. In our subsequent exploration, we integrated a self-attention module preceding the fully connected layer to discern its influence on model performance. The self-attention mechanism’s strength lies in its ability to capture long-range dependencies and discern correlations across various segments of the amino acid sequence. Nevertheless, our experimental results indicated that MetalPrognosis’s performance dipped noticeably upon integrating the self-attention module for single metal datasets, although there was a minor enhancement in MCC for the mixed dataset. We theorize that the increased computational complexity introduced by the self-attention module, combined with our training data’s limited size, might have led to overfitting. Furthermore, given the sufficient receptive field offered by the two-layer convolution model, including an attention module might not contribute additional benefits but augment the overall computational demands. In summation, our ablation studies affirm the logical design of the MetalPrognosis framework. The embedding extracted from its final layer effectively captures dynamic semantic information pertinent to protein structure and function.

### 3.3 Comparison with state-of-the-art methods on the independent test set

In order to verify the generalization ability of the MetalPrognosis on unknown data, we compared its prediction performance with two state-of-the-art machine learning methods on the independent test set. There are 118 disease-related metal binding sites in the independent test set, with 59 pathogenic mutation sites and 59 benign mutation sites respectively, including metals such as zinc, calcium, and magnesium. We combined the mixed data set with different metalloproteins and the benign mutation data set to train the model, then saved the model weight of the best training epoch for verification on the independent test set.

It can be seen from table 4 and figure 2B that the model surpasses the other two machine learning methods based on structural features and multiple sequence alignment features in several indicators. Specifically, it is 0.096, 0.089, 0.034, 0.022, 0.076, and 0.073 higher than the most advanced structure-based deep learning method MCCNN on specificity, precision, accuracy, F1-score, MCC, and AUC respectively. From the ROC curve in Figure 2C, it can be seen that the prediction performance of MetalPrognosis on the independent test set is much stronger than that of MCCNN. However, the sensitivity of our model is slightly lower than that of MCCNN, which may be caused by the lack of data in the independent test set. We will collect more mutation sites to train the model to improve this indicator. It proves that informative dynamic sequence embedding extracted from the language model outperforms the energy-based affinity grid maps as spatial features and traditional static manually-engineered amino acid sequence features for binding site detection. This alignment-free method is efficient and fast because it avoids the time and labor consumption of multiple sequence alignment algorithms. The results of the independent test set further demonstrated the robustness and generalization ability of our model.

**Table 4:** Performance comparison of MetalPrognosis with state-of-the-art methods on the independent test set.

|  | Sensitivity | Specificity | Precision | Accuracy | F1-score | MCC | AUC |
| --- | --- | --- | --- | --- | --- | --- | --- |
| MetalPrognosis | 0.764 | **0.943** | **0.929** | **0.856** | **0.839** | **0.721** | **0.950** |
| MCCNN | 0.796 | 0.847 | 0.840 | 0.822 | 0.817 | 0.645 | 0.877 |
| PolyPhen-2 | **0.931** | 0.610 | 0.701 | 0.769 | 0.800 | 0.570 | – |

### 3.4 Webserver development

An online tool can provide great convenience for biologists or medical researchers who are not familiar with computer science. Therefore, we developed and deployed a free online web server for the proposed method to predict disease-related mutation sites on metalloproteins at http://metalprognosis.unimelb-biotools.cloud.edu.au/. The web page of MetalPrognosis was developed based on PHP, and the webserver is managed by the Nginx server on a 4-core Linux server machine with 16 GB RAM and 200 GB hard disk. Users can choose to upload the metalloproteins FASTA sequence of Homo sapiens, or the protein PDB structure and the corresponding metal binding sites, and then click submit, the backend server will return the prediction results later. We also provided input examples of fasta sequences and PDB protein structures, respectively. The prediction results will be displayed in a table, which can be exported to TXT, CSV, JSON, and XML format. Besides, it only takes less than one minute to predict ten metal binding sites, which proves the efficiency of our method.

### 3.5 Case studies

In order to further verify the predictive performance of MetalPrognosis for disease-related mutation sites, we selected three proteins from the independent test set for a case study. The first protein is P50461(Uniprot ID: P50461, Gene Name: CSRP3, PDB ID: 2O13), which is shown in Figure 3A. Our method can predict that the mutation at the 58th amino acid site is a pathogenic mutation related to the disease, and the MCCNN predicts incorrectly. Specifically, this site is a zinc ion binding site, and mutations from Cysteine to Glycine can cause cardiopathy. A hereditary heart disorder characterized by ventricular hypertrophy, which is usually asymmetric and often involves the interventricular septum. The symptoms include dyspnea, syncope, collapse, palpitations, and chest pain[48,49,50]. Besides, it will decrease interaction with NRAP and ACTN2, decrease zinc-binding and impair protein stability, and decrease PKC/PRKCA activity[51]. The probability of disease-related mutation predicted by our model is 0.777, while the probability predicted by MCCNN is 0.26.

**Fig. 3.**
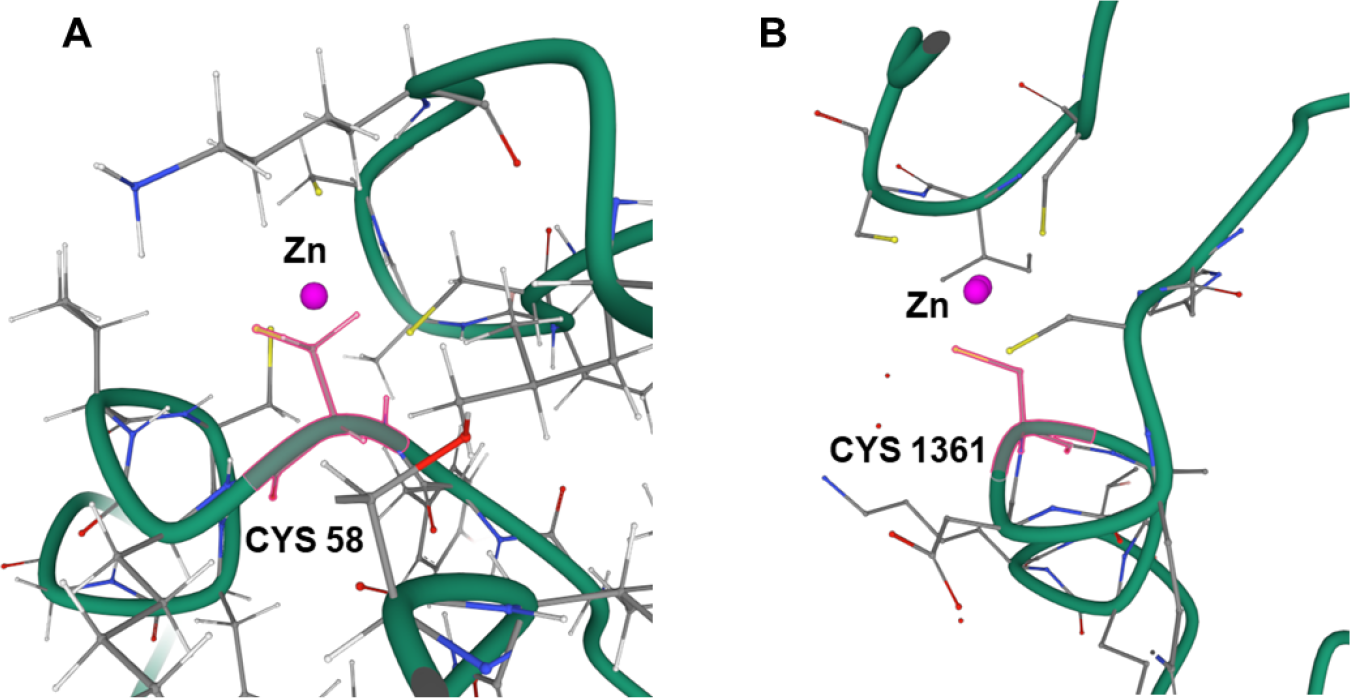
Visualization of two examples from the independent test set. **A**. Disease-associated mutation predicted by MetalPrognosis(Uniprot ID: P50461). **B**. Disease-associated mutation predicted by MetalPrognosis(Uniprot ID: O15550).

The second protein in Figure 3B is O15550(Uniprot ID: O15550, Gene Name: KDM6A, PDB ID: 3AVR). MetalPrognosis predicts that the mutation from cysteine to tyrosine at the 1361st site is a pathogenic mutation, with a prediction probability of 0.65. However, this site is annotated as a benign mutation by MCCNN, with a probability of 0.00004. This site is also a zinc ion binding site, and mutations can lead to a congenital intellectual disability syndrome with additional features, including postnatal dwarfism, a peculiar facies characterized by long palpebral fissures with eversion of the lateral third of the lower eyelids, radiographic abnormalities of the vertebrae, hands, and hip joints, and recurrent otitis media in infancy[52,53]. The last protein in Figure 2D is Q92730(Uniprot ID: Q92730, Gene Name: RND1, PDB ID: 2CLS). MetalPrognosis predicts that the mutation from proline to arginine on the 44th magnesium ion binding site is a benign mutation with a probability of 0.144. However, this site is annotated as a disease-related mutation by MCCNN with a probability of 0.783. In fact, the clinical significance of this site has not been reported in ClinVar.

## 4 Conclusion

Protein metal ion interactions play an important role in the pathogenesis of diseases. Missense mutations in protein metal binding sites can cause changes in the physiological environment and cause serious human diseases. However, current computational methods have limitations as they overlook dynamic semantic embeddings in amino acid contexts, posing a significant challenge for predicting pathogenic mutations at different metal sites In this work, we developed a framework called MetalPrognosis, which is a sequence-based alignment-free deep learning method for precisely predicting disease-associated mutation of metal-binding sites in metalloproteins. To capture long-term dependency information between amino acids, MetalPrognosis extracts dynamic semantic features from the last layer of the protein pre-trained language model ESM. After that, convolutional neural networks were introduced to obtain local information and integrate global abstract information. MetalPrognosis was shown to surpass state-of-the-art methods MCNN 0.096, 0.089, 0.034, 0.022, 0.073, and 0.076 in specificity, precision, accuracy, F1-score, MCC, and AUC respectively. In addition, the predicted results help to identify pathogenic missense mutation sites in human metalloproteins, demonstrating their potential for disease screening. We provide a publicly available online prediction platform and datasets at http://metalprognosis.unimelb-biotools.cloud.edu.au/ and https://github.com/Jrunchang/MetalPrognosis, respectively.

There is still room for further improvement in model performance. Due to the small number of training data samples used in this study, there are only disease-related mutations on the binding sites of three metals. The next step is to collect more data and build data sets of other metal ions. In the future, we will attempt to obtain protein tertiary structure and convert it to graph representations. Combining graph neural networks for more accurate identification of disease mutation sites.

