## Supplementary Material for "MetalPrognosis: a Biological Language Model-based Approach for Disease-Associated Mutations in Metal-Binding Site prediction"

### MetalPrognosis–Supplementary Material*

#### 1 Implementation details

We used five-fold cross validation strategy to train the model. It randomly divides the sample data into five parts, each time randomly selects four parts as the training set, and the remaining one part as the test set. It can better evaluate the generalization performance of the model and alleviate the overfitting. MetalPrognosis is developed based on PyTorch1.13.1, batch size is set to 16 and epoch is set to 100. We employed the Adam optimizer with a learning rate of 0.0001 and weight decay is 0.00001 to alleviate the overfitting[1]. Cross Entropy loss is used to reflect the difference between the ground truth and the predicted value of the model. The classification cross-entropy is defined as follows:

$$Loss = -\sum_{i=1}^{N} y_i \cdot log\hat{y}_i \tag{1}$$

where $\hat{y}_i$ is the predicted value, $y_i$ is the corresponding ground truth, and N is the 2 classes. The convolution kernel in the two-layer convolution neural network is 3 and the stride is 1. The kernel size and stride of max pooling layer are set to 2. In addition, by introducing activation function ReLU (Rectified Linear Unit) to improve the nonlinear expression ability of the network[2]. MetalPrognosis is trained on a powerful Linux server machine, which has a 32-core central processing unit (CPU), 2 NVIDIA GeForce RTX 3090 Ti graphics processing unit (GPU) and 128 GB RAM. High performance server provides strong support for model training and validation.

#### 2 Performance evaluation

In order to measure the predictive performance of MetalPrognosis, this study used evaluation metrics of common binary classification problems to evaluate the model. including accuracy, precision, recall, F1-score, specificity, Matthews correlation coefficient (MCC) and area under the receiver operating characteristic curve (AUC), which are defined below:

$$Accuracy = \frac{(TP + TN)}{TP + TN + FP + FN} \tag{2}$$

$$Recall = \frac{TP}{(TP + FN)} \tag{3}$$

$$Precision = \frac{TP}{(TP + FP)} \tag{4}$$

$$Specificity = \frac{TN}{(TN + FP)} \tag{5}$$

$$F1 = \frac{2 \times Precision \times Recall}{(Precision + Recall)} \tag{6}$$

$$MCC = \frac{(TP \times TN - FP \times FN)}{\sqrt{(TP + FP)(TP + FN)(TN + FP)(TN + FN)}} \tag{7}$$

Where TP, TN, FP, and FN represent the numbers of true positives, true negatives, false positives and false negatives, respectively. Accuracy refers to the proportion of correctly predicted samples to all samples. Recall, like sensitivity, is the proportion of the number of correctly predicted positive samples to all positive samples. Precision is the proportion

* Fuyi Li is the corresponding author, he is supported by the National Natural Scientific Foundation of China (No. 62202388), the National Key Research and Development Program of China (No. 2022YFF1000100), the Qin Chuangyuan Innovation and Entrepreneurship Talent Project (No. QCYRCXM-2022-230), and Talent Research Funding at Northwest A&F University (No. Z1090222021). The first two authors contributed equally to this study.

of the number of correctly predicted positive samples to the number of all predicted positive samples. Besides, F1-score and MCC are more suitable for measuring situations where data is imbalanced, which can more comprehensively measure the performance of the model. Additionally, we also plotted ROC curves to visualize the trade-off between true positives and false positives.

#### 3 Characterization of disease-associated mutations at metal-binding sites

Previous statistical in MCCNN results showed that most disease-related amino acids in metal binding sites had a high frequency of mutation from neutral to hydrophilic and from neutral to hydrophobic, accounting for 28% and 24% respectively. Besides, a large number of disease associated mutations lead to characteristic nonpolar to polar changes[3]. According to statistics, metal proteins with the same type of metal elements at multiple sites have the highest proportion, accounting for 79.1%[4]. Figure 2 shows the quantitative distribution of amino acids of disease-related mutations at three different metal sites.

It is obvious that the amount of cysteine is the largest in zinc ions in Figure 1A, more than 100, followed by arginine, histidine and glycine. On the missense mutation of calcium binding site(Figure 2B), the amount of aspartic acid was the largest, close to 80, followed by cysteine and glycine. It can be seen from the figure 1 that the number of mutations in cysteine and glycine is high in both metalloproteins. Similarly, at the magnesium ion binding site in Figure 1C, the maximum number of glycine is 30. But the number of magnesium ion metalloprotein is the least, only 171. Although the data sample on the independent test set is small, the number of cysteine in disease-related mutations is 17, more than a quarter of the total.

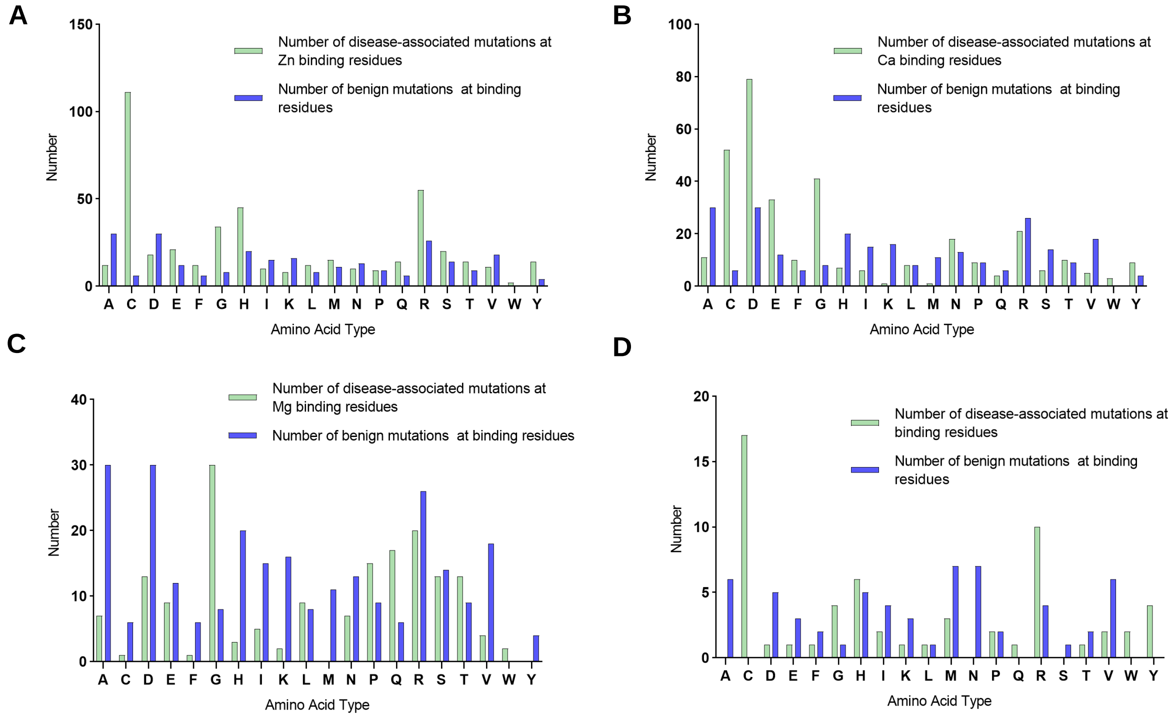


**Fig. 1.** Columnar distribution of amino acids on three metal datasets. **A.** Amino acid distribution at zinc ion binding sites. **B.** Amino acid distribution at calcium ion binding sites. **C.** Amino acid distribution at magnesium ion binding sites. **D.** Amino acid distribution on the independent test set.
